# REST-AKT-SOX2 axis controls neuronal lineage state and Midkine-mediated communication in SHH-medulloblastoma

**DOI:** 10.64898/2026.04.06.716597

**Authors:** Ashutosh Singh, Lei Guo, Jyothishmathi Swaminathan, Donghang Cheng, Yanwen Yang, Swarnalatha Manickavinayaham, Lin Xu, Vidya Gopalakrishnan

## Abstract

Stem/progenitor-like cells are known drivers of tumor progression, chemoresistance, and relapse in Sonic-Hedgehog medulloblastoma (SHH-MB), yet the regulatory mechanisms that sustain these resilient cellular states remain incompletely defined. Here, we identify the *RE1*-silencing transcription factor (REST) as a key transcriptional regulator that preserves the progenitor compartment in SHH-MB through stabilization of the stemness factor SOX2. Mechanistically, REST activates AKT signaling, which in turn enhances SOX2 protein stability, revealing a REST-AKT-SOX2 axis that supports stem/progenitor identity and ongoing tumor maintenance. Beyond maintaining intrinsic stem-like programs, REST also orchestrates the extrinsic communication network of SHH-MB tumors. Single cell transcriptomic profiling and ligand-receptor interaction mapping highlight Midkine (MDK)-mediated signaling as one of the most upregulated intercellular communication routes in MB. We demonstrate that REST drives this signaling cascade through its control of SOX2. Perturbation of REST or SOX2 results in reduced MDK and its receptor, SDC2 expression, and chromatin immunoprecipitation assays show that SOX2 directly binds and regulates *MDK*/*SDC2* expression, establishing and reinforcing a REST/SOX2 centered transcriptional mechanism that coordinates both progenitor maintenance and cell-cell communication in the malignant compartment. Together, these findings position REST as an integrator of intrinsic progenitor cell programs and extrinsic MDK-mediated signaling in SHH-MB. By linking stemness, communication and potential for therapeutic resistance, the REST-AKT-SOX2-MDK signaling axis emerges as a targetable vulnerability to suppress stem/progenitor driven tumor program in REST-driven SHH-MBs.

## Introduction

Medulloblastoma (MB) is the most common malignant pediatric brain tumor and accounts for approximately 20% of childhood central nervous system cancers in the United States [1–4]. Despite significant advances in multimodal therapy, MB remains a leading cause of cancer-related morbidity and mortality in children [5, 6]. High throughput genomic and transcriptomic analyses have established that MB comprises four principal molecular subgroups - WNT, Sonic Hedgehog (SHH), Group 3, and Group 4, each arising from distinct developmental lineages and defined by characteristic genetic alterations and clinical behaviors [7–9]. This refined molecular taxonomy has transformed therapeutic strategies, enabling risk-adapted treatment approaches that combine maximal safe resection, craniospinal irradiation, and multi-agent chemotherapy [10, 11]. While patients with WNT-driven tumors achieve survival rates exceeding 90%, outcomes remain suboptimal for children with SHH and Group 4 tumors, with the worst outcomes seen in patients with Group 3 tumors, underscoring the urgent need for more effective and biologically informed interventions [6, 11].

A major obstacle to improving survival is the high incidence of treatment resistance and tumor recurrence, which occurs in nearly one-third of patients and is strongly associated with poor prognosis [11, 12]. Growing evidence suggests that a population of stem/progenitor-like tumor cells underlies this therapeutic failure [13, 14]. These resilient cells frequently exhibit enhanced DNA damage repair, metabolic adaptability, and intrinsic tolerance to cytotoxic stress, enabling them to withstand initial therapy and subsequently regenerate recurrent tumors [15, 16]. Defining the molecular regulators that sustain such progenitor-like states is therefore critical to understanding mechanisms of persistence and identifying new therapeutic vulnerabilities.

One compelling candidate is the *RE1*-silencing transcription factor (REST), a master regulator of neural progenitor identity and suppressor of neuronal differentiation [17, 18]. We previously showed *REST* gene expression is aberrantly elevated in SHH MBs, which is associated with increased risk for metastasis and poor survival [19–21]. Previous studies have also implicated REST in maintaining an undifferentiated, progenitor-like phenotype and promoting tumor cell survival, suggesting that REST may have the potential to be a driver of treatment resistance and disease relapse [22, 23]. Given its dual functions in lineage control and tumor maintenance, REST could be therapeutically target. However, transcription factors are difficult to target, highlighting the need to better understand downstream mechanisms through which it sustains stem/progenitor-like states in MB to identify other druggable opportunities.

Cell-cell communication in cancers is often hijacked by malignant cells, enabling them to evade normal regulatory controls and promoting uncontrolled growth, survival, and metastasis [24, 25]. Cancer cells disrupt both direct cell-cell contact and secreted signaling pathways to reprogram the tumor microenvironment in ways that support tumor progression and therapy resistance [26, 27]. MB is no exception; it relies heavily on dysregulated signaling mechanisms such as Midkine (MDK) and Reelin (RLN)-mediated pathways to sustain tumor growth and drive chemoresistance [28–30]. These aberrant signaling cues are known to promote stem cell-like transitions within the tumor microenvironment by stimulating the proliferation of quiescent neuronal progenitor cells, the primary cells of origin in SHH-MB, ultimately contributing to tumor recurrence [31–33]. However, regulators of these aberrant signaling circuits remain to be fully delineated.

In this study, we identify REST as a central regulator linking intrinsic progenitor maintenance to extrinsic cell-cell communication in SHH-MB. We show that REST stabilizes the stemness factor SOX2 through activation of AKT signaling, establishing a REST-AKT-SOX2 axis that preserves stem/progenitor-like cell populations and supports tumor maintenance. We further demonstrate that this axis drives elevated MDK-mediated signaling, one of the most prominent communication routes in MBs, by promoting SOX2-dependent transcription of *MDK* and its receptor/co-receptor partner - SDC2. Although enhanced MDK signaling has been previously observed in MBs, mechanisms underlying this phenomenon are unexplained. Our findings provide one of the first mechanistic explanation for elevated MDK-SDC2 and MDK-NCL signaling observed in SHH-MB and establish REST as a key integrator of stemness and intercellular communication programs that may fuel tumor persistence and relapse.

## Materials and Methods

### Patient samples

Two publicly available MB transcriptomic datasets (GSE85217 and GSE148389) were analyzed to assess gene expression patterns [9, 34]. Differential expression analyses were conducted using the R2 Genomics Analysis and Visualization Platform (http://r2.amc.nl). Statistical significance was defined as p < 0.05. Gene ontology p-values are not corrected for multiple testing.

### Single cell RNA seq analysis

Single-cell RNA-sequencing (scRNA-seq) data from pediatric MB samples were obtained from the Gene Expression Omnibus database under accession number GSE155446 [35] (comprising samples from 28 patients across the four main molecular subgroups: WNT, SHH, Group 3, and Group 4). Raw count matrices were processed and analyzed using the Seurat in R. Following quality control, normalization, and identification of highly variable features, datasets from multiple samples were integrated using Seurat’s integration workflow to correct for batch effects. Dimensionality reduction was performed via principal component analysis (PCA), followed by graph-based clustering and visualization using Uniform Manifold Approximation and Projection (UMAP). Major cell types, including cerebellar granule neural precursors (CGNPs), were annotated based on established canonical marker genes. Within the integrated dataset, cells were stratified into two groups- Low-REST and High-REST, based on whether expression of the REST gene was below or above its median expression level across all cells. CGNPs were subsetted from the integrated object and subjected to re-clustering using Seurat to resolve finer subpopulations. Cell-cycle phase and other cell types were identified using a predefined set of marker genes.

Cell-cell communication within the CGNP subpopulation and between Low-REST and High-REST CGNPs was inferred using the CellChat package in R [36]. Ligand-receptor interactions were predicted based on expression patterns, with default parameters unless otherwise specified. Pseudotime trajectory analysis of the CGNP subpopulation was conducted using Monocle 3 in R to infer differentiation trajectories and branching points [37]. All analyses were performed in R 4.3, with package versions documented in the session information (available upon request).

### Cell culture

Four SHH-MB cell lines, DAOY, UW228, UW426, and ONS76 were used in this study. The UW228 and UW426 lines were generously provided by Dr. John Silber (University of Washington). DAOY cells were obtained from the American Type Culture Collection (ATCC, Manassas, VA), and ONS76 cells were purchased from Accegen (New Jersey, USA). All the lines were maintained in Dulbecco’s modified Eagle’s medium (Sigma-Aldrich, MO, USA), supplemented with 10% fetal bovine serum (Sigma-Aldrich), 1% antibiotic-antimycotic (Thermo Fisher Scientific, MA, USA), and 1% sodium pyruvate (Thermo Fisher Scientific) and grown at 37 °C with 5% CO2.

### Patient-derived xenograft models

Patient-derived orthotopic xenograft (PDOX) models (RCMB-18, RCMB-24, and RCMB-54) were generously provided by Dr. Robert Wechsler-Reya (Columbia University). Tumors were serially propagated in NOD.Cg-*Prkdc^scid^ Il2rg^tm1Wjl^*/SzJ (NSG) mice (Jackson Laboratory, Bar Harbor, ME), through intracranial implantation using a stereotactic apparatus, following previously described procedures [20]. All mouse housing, care, and experimental procedures were performed under a protocol approved by the Institutional Animal Care and Use Committee (IACUC) of The University of Texas MD Anderson Cancer Center. Reporting of animal experiments adheres to ARRIVE guidelines (https://arriveguidelines.org).

### Transgenic animals

The generation of *Ptch^+/−^/REST^TG^* transgenic mice is previously described by Dobson et al [20]. The hREST transgene expression was induced through intraperitoneal administration of tamoxifen (100 µL of a 2 mg/mL solution; Sigma-Aldrich, Cat# T5648) on postnatal days 2, 3, and 4. Animals exhibiting moribund behavior were humanely euthanized, and tumor-bearing brains were collected for subsequent analyses. All procedures involving animals were conducted under protocols approved by the institutional IACUC and were carried out in accordance with ARRIVE reporting standards. (https://arriveguidelines.org).

### Immunohistochemistry

Mouse brains were fixed in 10% buffered formalin phosphate for 48 h and subsequently processed for paraffin embedding. Tissue sections of 5 µm thickness were prepared and deparaffinized using a Gemini AS Automated Stainer (Thermo Fisher Scientific). Following overnight incubation with primary antibodies at 4 °C, sections were treated with biotinylated secondary antibodies supplied with either the ABC kit or the MOM kit (Vector Laboratories, CA, USA). Signal detection was performed using the VECTASTAIN® Elite® ABC-HRP Kit, Peroxidase (Vector Laboratories, Cat# PK-6101), according to the manufacturer’s protocol, and visualized with the DAB Peroxidase Substrate Kit (Vector Laboratories, Cat# SK-4100). Slides were then counterstained with hematoxylin, mounted, and examined using a Nikon ECLIPSE E200 microscope equipped with an Olympus SC100 camera. A complete list of primary antibodies employed for immunohistochemistry is provided in Table S1.

### Lentiviral transduction and transient transfection

Human embryonic kidney 293T (HEK293T) cells were co-transfected with either a control construct or the gene of interest (shRNA constructs) with the packaging plasmid psPAX2 and the envelope plasmid MD2. Lentiviral supernatants were harvested 48 hours after transfection and the lentivirus were purified by 0.45 μm PES filter. MB cells were subsequently transduced with the collected viral medium supplemented with Polybrene (8 μg/mL) and allowed to incubate for 48 hours. Following transduction, cells were placed under selection with puromycin (2 μg/mL) for up to one week. For transient knockdown of SOX2, cells were transfected for 48 hours with MISSION® esiRNA targeting human SOX2 (Sigma Aldrich, Cat# EHU184131) or with the MISSION® Universal Negative Control siRNA (Sigma Aldrich, Cat# SIC001) using Lipofectamine 3000 (Thermo Fisher Scientific, Cat# L3000015).

### Western blot analyses

Cell lysates were prepared using lysis buffer composed of 50 mM Tris-HCl (pH 8.0), 50 mM NaCl, 1% NP-40, 0.5% sodium deoxycholate, 0.1% SDS, and protease/phosphatase inhibitor cocktail. Protein samples were processed for Western blotting following previously described procedures [19]. Membranes were incubated with primary antibodies listed in Table S1 and subsequently with HRP-conjugated goat anti-mouse or anti-rabbit secondary antibodies. Chemiluminescent detection was performed using either the SuperSignal West Dura Extended Duration Substrate (Thermo Fisher Scientific, Cat# 34075) or the Western Lightning Plus-ECL Enhanced Chemiluminescence Substrate (Fisher Scientific, Cat# 50-904-9325). Blots were visualized using a ChemiDoc Touch Imaging System (Bio-Rad). Image quantification and analysis were conducted with ImageJ software.

### Co-immunoprecipitation

Cell pellets were washed with ice-cold PBS and subsequently lysed in a mild lysis buffer composed of 50 mM Tris-HCl (pH 7.5), 150 mM NaCl, 1 mM EDTA, 1% Triton X-100, and 5 mM EDTA. The buffer was supplemented with a Protease/Phosphatase Inhibitor Cocktail (100X) (Cell Signaling Technology, Cat# 5872S). The samples were then sonicated to ensure thorough lysis. For immunoprecipitation, clarified lysates were incubated overnight at 4°C with either control mouse IgG or primary antibodies, anti-UBR5 (Cell Signaling Technology, Cat# 65344). Immune complexes were captured by incubation with Pierce™ Protein A/G UltraLink™ Resin (Thermo Fisher Scientific, Cat# 53132) for 1.5 hours at 4°C. Following four washes with lysis buffer, the bound proteins were eluted by boiling the beads in loading buffer. Samples were resolved by SDS-PAGE, transferred onto PVDF membranes, and subsequently analyzed by Western blotting.

### Quantitative real-time PCR

Total RNA was extracted using the Quick-RNA MiniPrep Kit (Zymo Research, Cat# R1055). For cDNA synthesis, 1 μg of RNA was reverse-transcribed using the iScript cDNA Synthesis Kit (Bio-Rad, Hercules, CA). Quantitative real-time PCR (qRT-PCR) was conducted in triplicate with the 2X SensiMix SYBR & Fluorescein Kit (Bioline, Boston, MA) on a LightCycler 96 Real-Time PCR System (Roche Diagnostics GmbH, Mannheim, Germany). Relative gene expression levels were quantified using the comparative 2^−ΔΔCp method, with normalization to 18S ribosomal RNA, and results are presented as fold change relative to controls. Primer sequences used in this study are provided in Table S2.

### Chromatin Immunoprecipitation

The SHH-MB cell lines ONS76 and DAOY were fixed with 1% formaldehyde to induce cross-linking and subsequently processed for chromatin immunoprecipitation (ChIP) following previously established procedures [20]. In brief, cross-linked cells were washed with 1× phosphate-buffered saline (PBS) and lysed in a buffer containing 50 mM Tris-HCl (pH 8.0), 10 mM EDTA (pH 8.0), 1% SDS, and protease inhibitors. The lysates were then sonicated to shear chromatin, and 1% of the resulting material was retained as input DNA. The remaining chromatin was diluted in ChIP dilution buffer (16.7 mM Tris-HCl, pH 8.0; 167 mM NaCl; 1.2 mM EDTA, pH 8.0; 1.1% Triton X-100; plus protease inhibitors), precleared, and incubated with the SOX2 monoclonal antibody (Thermo Fisher Scientific, Cat# MA1-014) for 12 hours at 4°C. Immune complexes were captured using protein A beads (Millipore), followed by sequential washes and elution. After reversal of cross-links, DNA was purified using a PCR purification kit (Zymo Research). Quantification of immunoprecipitated DNA was performed using SYBR Green-based qPCR on a Roche LightCycler 96 instrument, and relative enrichment was calculated using the comparative 2^−ΔΔCp method. Primer sequences used in these assays are listed in Table S2.

### Drug studies

To assess the effect of capivasertib, ipatasertib, and vevorisertib (MedChem Express, Cat# HY-15431, HY-15186, and HY-137458, respectively), on MB cell viability, cells were plated in 96-well plates at a density of 5 × 10³ cells per well. Cells were then treated with vehicle control (0.2% DMSO) or increasing concentrations of Capivasertib for 48 hours. Cell viability was quantified using the MTT assay, and absorbance was recorded at 570 nm using the CLARIOstar Plus plate reader.

### Statistical analysis

Statistical analyses were performed using GraphPad Prism version 10. Data are presented as mean ± standard deviation (SD), derived from at least three independent biological replicates. Comparisons of means between groups were conducted using unpaired Student’s t tests. To analyze transcriptomic data from MB patients, Welch’s t test was used to account for unequal variances and sample sizes. Statistical significance was defined as p < 0.05, and results are denoted as: ****p < 0.0001, ***p = 0.0001-0.001, **p = 0.001-0.01, *p = 0.01-0.05, ns = not significant.

## Results

### REST elevation is associated with proliferating CGNP-enriched clusters

Studies from our group, and others have shown that REST elevation contributes to MB-genesis [20, 38]. To better define key pathways affected by REST elevation, comparative pathway enrichment analyses of high-REST and low-REST SHH-MBs were performed on two independent publicly available transcriptomic datasets, GSE85217 and GSE148389 (Fig. 1A). As expected, pathways related to chromatin organization and remodeling, cell-cycle, neurogenesis, and neuronal differentiation were significantly enriched. Additionally, pathways related to cell migration, hypoxia, angiogenesis, autophagy, and ubiquitination were also enriched, findings consistent with recent studies from our group [19, 21, 39, 40]. Interestingly, stemness-related pathways, including stem cell proliferation, development and maintenance, and cerebellar granule neural precursor (CGNP) proliferation were highly represented in high-REST SHH-MBs compared to low-REST samples (Fig. 1A).

**Figure 1:**
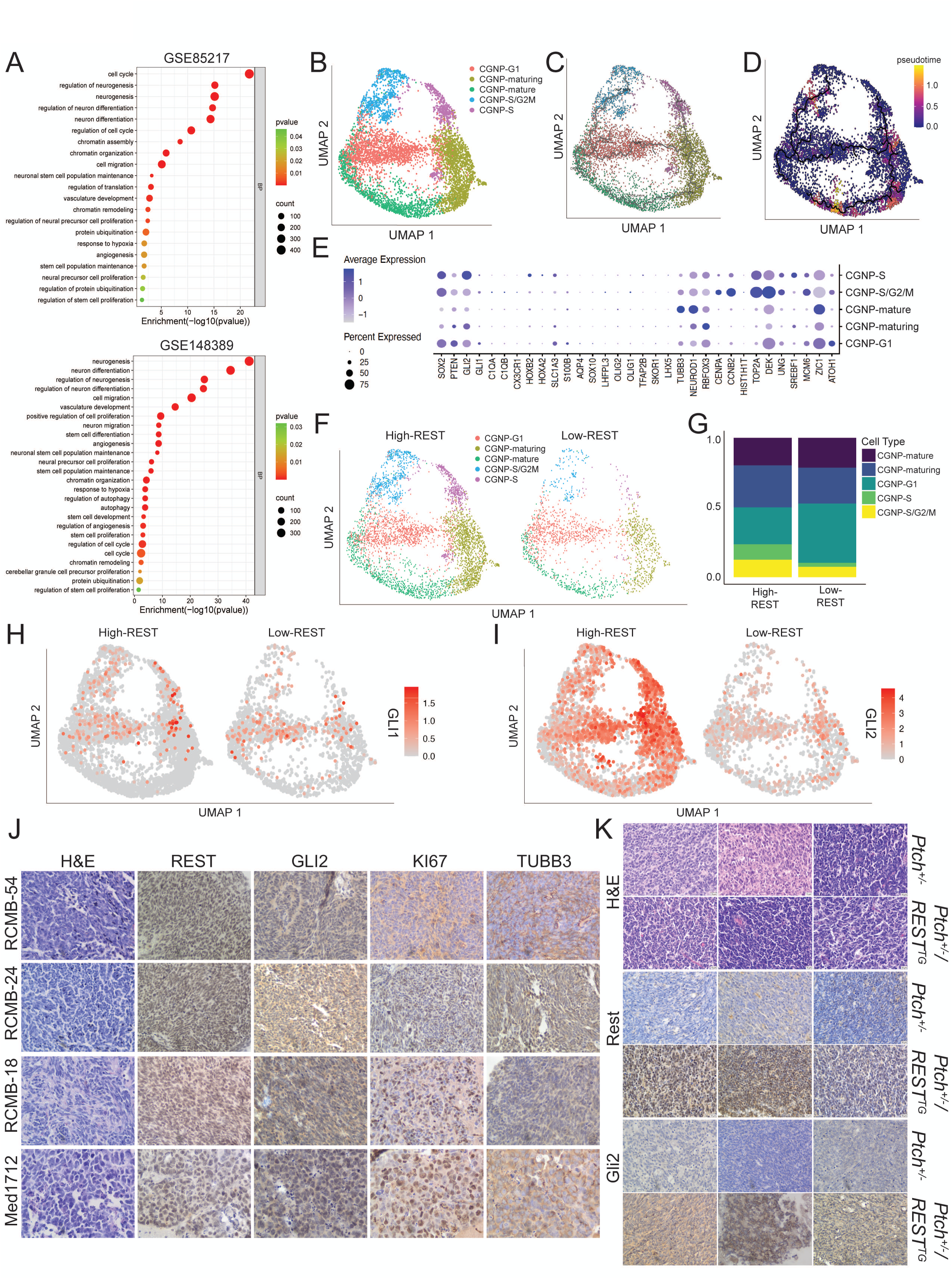
REST expression in SHH-MB tumors is associated with increased population of premature cells exhibiting high GLI2 expression. (A) Bubble plot showing enriched Gene Ontology (GO) terms in high-REST compared to low-REST SHH-MB samples in two transcriptomic datasets, GSE85217 and GSE148389. (B) UMAP plot of scRNAseq data (GSE155446) of SHH-MBs illustrating distinct transcriptional and neuronal hierarchical states, with each cluster representing a CGNP-related subpopulation- CGNP-G1, CGNP-S, CGNP-S/G2M, and CGNP-Maturing and CGNP-Mature. (C) UMAP plot showing cell trajectory, highlighting transcriptional transitions from CGNP proliferation to maturation states, with progression inferred by color gradients. (D) UMAP plot overlaid with pseudotime analysis, illustrating the continuous developmental trajectory from proliferating CGNPs toward more differentiated cell states. (E) Dot plot showing the expression of key marker gene signatures that define the CGNP-G1, CGNP-S, CGNP-S/G2M, and CGNP-Maturing and CGNP-Mature. subpopulations in GSE155446 SHH-MB scRNAseq dataset. Dot size represents the percentage of cells expressing each gene within a given cluster, while color intensity reflects the average expression level. (F) UMAP plot showing the differences in the above cell subclusters following stratification of the SHH-MB scRNAseq dataset into high-REST and low-REST groups. (G) Stacked bar plot quantifying the proportional distribution of different CGNP clusters within the high- and low-REST cohorts. UMAP plot of (H) *GLI1* and (I) *GLI2* expression in the CGNP clusters from F. IHC of cerebellar sections from (J) mice with SHH-MB PDOX and (K) tumor-bearing *Ptch^+/−^* and *Ptch^+/−^/REST^TG^* animals (n = 3) for REST, Ki67, TUBB3, GLI2, and Rest and Gli2 respectively (400X magnification).

These findings were further validated through analysis of the GSE155446 single-cell RNA-seq dataset (Fig. 1B). Focusing primarily on the malignant compartment in SHH-MB samples, unsupervised clustering and UMAP visualization revealed distinct populations corresponding to CGNPs at different cell-cycle stages (G1, S, and S/G2M), maturing and mature CGNPs (Fig. 1B), indicating heterogeneity within the tumor. We then performed trajectory analysis to define lineage relationships and demonstrate a continuous progression from cycling CGNPs toward maturing and matured CGNPs (Fig. 1C). Pseudotime ordering further supported this developmental hierarchy, positioning proliferative CGNPs at the early point of the trajectory and mature CGNPs at the terminal stage of the path (Fig. 1D). Transcriptomic profiling of these malignant subclusters indicated that CGNPs marked by *ZIC1* expression were the predominant cells in the population highlighting the role of neural precursors in SHH-MB genesis (Fig. 1E). Minimal expression of marker genes for GABAergic neurons (*LHX5*, *SKOR1*, and *TFAP2B*), glutamatergic neurons (*HOXA2* and *HOXB2*), oligodendrocytes (*OLIG1*, *OLIG2*, *LHFPL3*, and *SOX10*), astrocytes (*AQP4*, *S100B*, and *SLC1A3*), and immune cells (*CX3CR1*, *C1QB*, and *C1QA*) confirmed that the analyzed samples consisted of population selected for malignant cells (Fig. 1E). All *ZIC1*-expressing CGNPs were stratified into two subpopulations: proliferative CGNPs (*ATOH1* positive), and postmitotic CGNPs (*RBFOX3*, *NEUROD1*, and *TUBB3*). Surprisingly, strong *GLI2* but not *GLI1* expression was seen in three *CGNP* clusters corresponding to cell-cycle phases: G1 (*MCM6*), S (*DEK* and *TOP2A*), and S/G2M (*DEK*, *TOP2A*, *CENPA*, and *CCNB2*) (Fig. 1E). Additionally, these clusters were also associated with robust *SOX2* expression (Fig. 1E).

Next, to follow up on REST-dependent enrichment of processes involved in stemness, and to assess if REST levels influenced these cell states, we stratified *ZIC1+* malignant cells into high-REST and low-REST groups and visualized their distribution across the UMAP (Fig. 1F and 1G). As shown in Fig. 1F, when compared to the low-REST cluster, high-REST cluster contained a larger proportion of proliferative CGNPs spanning three cell-cycle stages, along with mature CGNPs. The high-REST population was primarily associated with CGNPs in the G1, S and S/G2M phases, whereas the low-REST samples consisted mainly of CGNP-G1 and CGNP-Mature clusters (Fig. 1G). These patterns suggest that higher *REST* expression is associated with CGNP immaturity and proliferative capacity while lower REST expression is aligned with neuronal maturation, consistent with its role as negative regulator of cell-cycle exit and terminal neuronal differentiation of CGNPs [17, 41].

### REST expression correlates with GLI2 expression in proliferating immature CGNPs

GLI transcription factors are key mediators of SHH signaling and are central to SHH-MB development. Unsupervised analyses of the high- and low-REST malignant clusters showed that *GLI1* expression was limited to proliferating CGNP cell clusters in both high-and low-REST SHH-MBs (Figs. 1H). However, *GLI2* was robustly expressed in proliferating cyclic and maturing CGNPs in high-REST SHH-MBs compared to low-REST tumors (Fig. 1I). These observations were further confirmed by immunopathological analyses of brain sections from mice bearing PDOX of SHH-MBs where all samples exhibited REST expression, accompanied by positive staining for Ki67 and low TUBB3, confirming their neuronal origins. Notably, these sections also demonstrated elevated GLI2 levels (Fig. 1J). Cerebellar sections of SHH-MBs from transgenic mice with Rest elevation (*Ptch^+/−^/REST^TG^*) also showed strong expression of Gli2 (Fig. 1K). These data suggest that upregulation of REST expression is associated with an increase in GLI2 levels.

### REST elevation is associated with increased SOX2 levels and expression of its target genes

GLI2 is an established transcriptional regulator of *SOX2* [42], and *GLI2* and *SOX2* expression levels show a strong positive correlation in SHH-MB patient transcriptomic datasets (GSE85217) (r=0.372; p<0.0001) (Fig. 2A). siRNA-mediated knockdown of *GLI2* resulted in reduced *SOX2* expression in DAOY cells (Fig. S1). In addition, transcriptomic analyses of patient samples (GSE85217) revealed a positive correlation between *REST* and *SOX2* expression (r=0.258; p<0.0001) (Fig. 2B). This relationship was further validated in scRNAseq data of SHH-MB patients, where high-REST tumor cell clusters exhibited increased *SOX2* expression compared with low-REST clusters (Fig. 2C). Moreover, differential gene expression analysis across subclusters of high- and low-REST tumors demonstrated elevated expression of *GLI2*, *SOX2*, and multiple known SOX2 target genes including, *MYCN*, *SMC4*, and *EGR1* [43] specifically within proliferating and maturing CGNP populations of high-REST tumors (Fig. 2D). Immunohistochemical (IHC) staining also confirmed an association between REST and SOX2 expression in SHH-MB PDOX tumors (Fig. 2E). Further, higher Sox2 expression was observed in tumor sections of *Ptch^+/−^/REST^TG^* transgenic mice compared to *Ptch^+/−^* mice, confirming a REST-dependency on the changes in SOX2 levels in SHH-MBs (Fig. 2F). Western blot analysis of SHH-MB cell lines, ONS76 and DAOY, revealed higher REST and SOX2 levels compared to UW228 and UW426 cells, where both proteins were expressed at lower levels (Fig. 2G). Silencing of *REST* expression in ONS76 and DAOY cells led to a significant reduction in SOX2 levels (Fig. 2H). Conversely, ectopic REST promoted an increase in SOX2 levels (Fig. 2I). These findings suggest that REST drives SOX2 protein expression in SHH-MBs.

**Figure 2:**
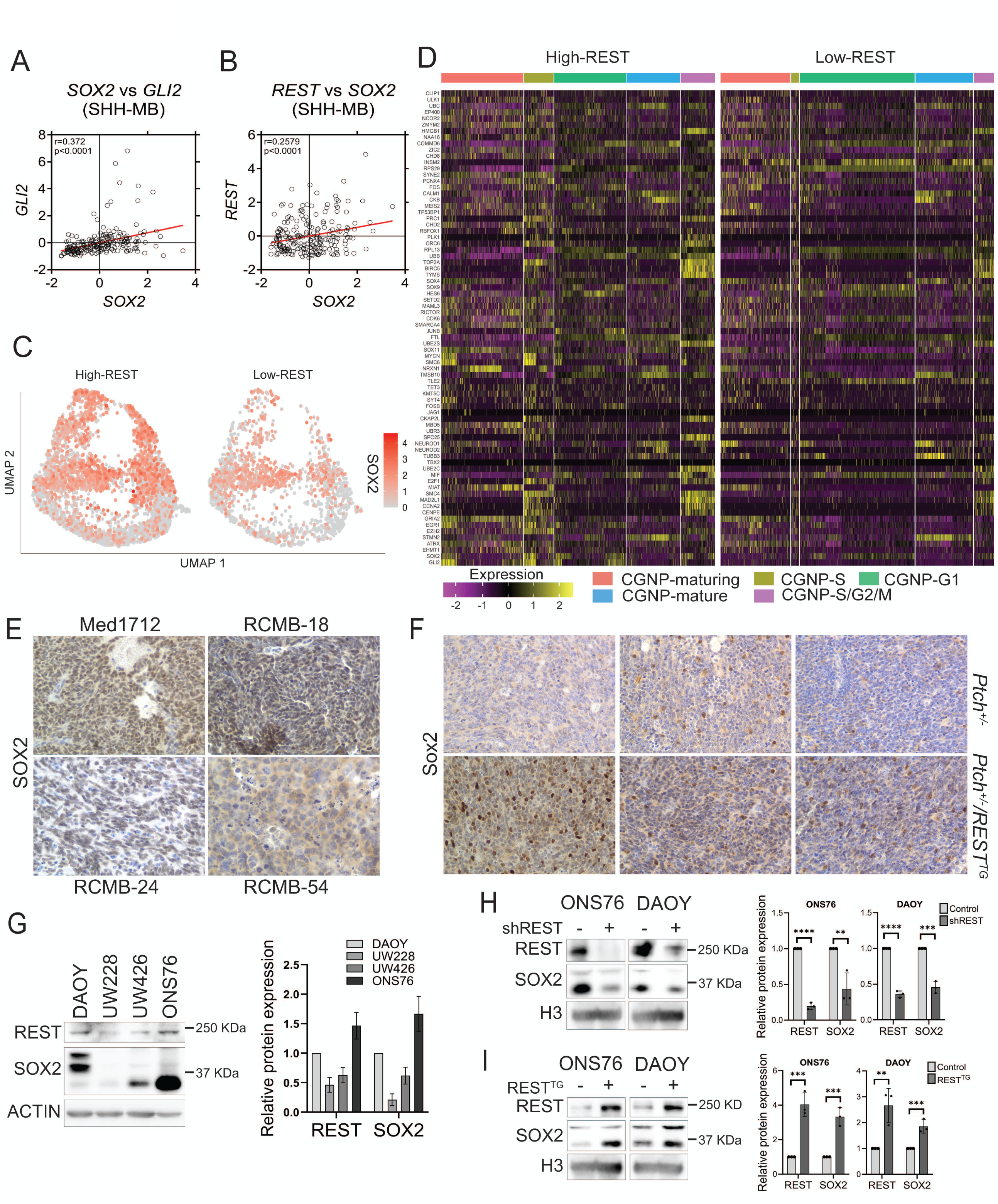
REST and SOX2 expression are correlated in SHH-MBs. Scatter plot to demonstrate correlation between **(A)** *SOX2* with *GLI2* and **(B)** *REST* with *SOX2* expression in GSE85217 microarray dataset (n = 223). **(C)** UMAP plot of *SOX2* expression in the CGNP clusters in 1F. **(D)** Heatmap to show expression differences of known SOX2-target genes in different CGNP clusters (G1, S, S/G2M, Maturing and Mature) in high- and low-REST SHH-MBs included in GSE155446. IHC to study SOX2 levels in cerebellar sections from **(E)** mice with SHH-MB PDOX and **(F)** tumor-bearing *Ptch^+/−^* and *Ptch^+/−^/REST^TG^* animals (n = 3) (400X magnification). Western blot to show **(G)** REST and SOX2 levels in DAOY, UW228, UW426, and ONS76 cells and change in their levels following **(H)** *REST* knockdown and **(I)** its overexpression in DAOY and ONS76 cells. Statistical data are presented for three independent biological replicates as the means ± SDs. **p < 0.01, ***p < 0.001, and ****p < 0.0001 by Student’s t test.

### REST-dependent SOX2 elevation is mediated by AKT-dependent protein stabilization

In previous work by Dobson et al [20], we reported that *REST* elevation in SHH-MBs is correlated with reduced *PTEN* expression, a negative regulator of AKT signaling. It is also known that AKT-dependent phosphorylation of SOX2 at threonine 116 (T116) residue, prevents its ubiquitination at the adjacent lysine 115 (K115) residue by the E3 ligase-UBR5, resulting in a blockade of proteasomal degradation of SOX2 [44] (Fig. 3A). scRNAseq analysis of SHH-MBs revealed *PTEN* expression was lower in most CGNP subclusters in high-REST SHH-MBs relative to the same subclusters in low-REST tumors (Fig. 3B). IHC of high-REST PDOX samples revealed low PTEN staining and strong p-AKT (Ser473) positivity and recapitulated previous findings in brain sections of tumor bearing *Ptch^+/−^/REST^TG^* and *Ptch^+/−^* mice by Dobson et al [20] (Fig. 3C). These observations confirm that REST elevation is associated with AKT activation.

**Figure 3:**
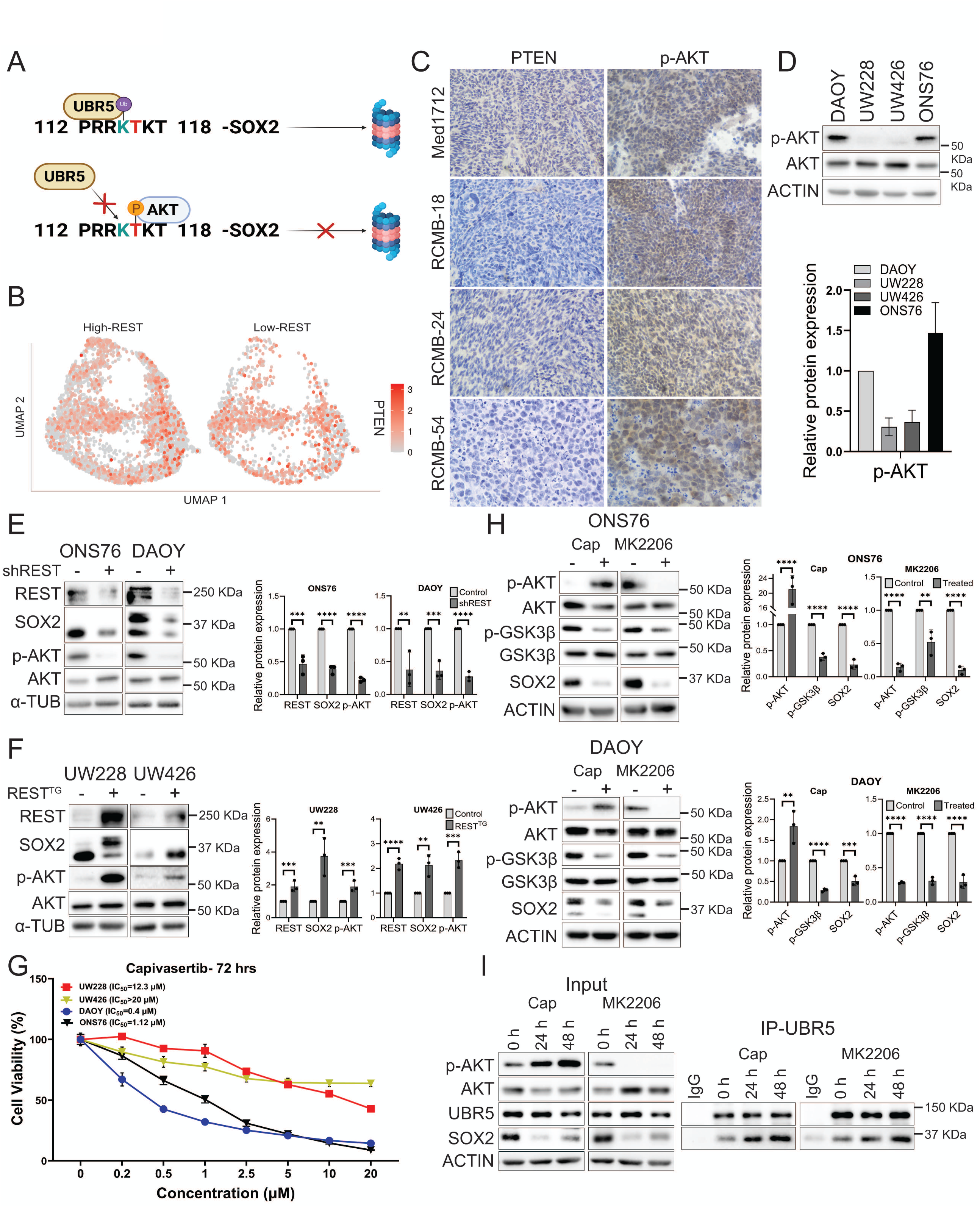
AKT activity is required for REST-dependent SOX2 stabilization in SHH-MB cells. **(A)** UBR5-mediated ubiquitination of SOX2 at K115 in the PRRKTKT motif (aa 112-118) targets it for proteasomal degradation. Phosphorylation of the adjacent residue T116 in SOX2 by AKT interferes with UBR5-dependent ubiquitination and degradation. **(B)** UMAP plot to show expression of *PTEN* in the CGNP clusters of high- and low-REST SHH-MB samples from 1F. **(C)** IHC for PTEN, p-AKT in cerebellar sections from mice with SHH-MB PDOX (400X magnification). **(D)** Western blot analysis to show basal level of phospho-(Ser473) and total AKT levels in DAOY, UW228, UW426, and ONS76 cells. A change in levels of REST, SOX2, p-AKT (Ser473), and AKT following REST knockdown **(E)** and overexpression **(F)** in DAOY/ONS76 cells, and UW228/UW426 cells, respectively, was measured through Western blot. α-tubulin was used as a loading control. **(G)** MTT assays to show sensitivity of DAOY, UW228, UW426, and ONS76 to various doses of capivasertib after 72-hour of drug treatment. IC_50_ values are included in the inset. **(H)** Western blot analysis of ONS76 and DAOY cells to show the effect of AKT inhibition by Capivasertib and MK2206 on p-AKT (Ser473), pGSK3β (Ser9), and SOX2 levels. Total AKT and GSK3β and Actin were included as controls. **(I)** Co-immunoprecipitation assay to show increased UBR5-SOX2 interaction and SOX2 destabilization following treatment with Capivasertib and MK2206. Anti-UBR5 or control IgG were used for pull-down. Western blots were performed for the indicated proteins. Actin served as a loading control. Statistical data are presented for three independent biological replicates as the means ± SDs.s **p < 0.01, ***p < 0.001, and ****p < 0.0001 by Student’s t test.

To assess if REST-dependent AKT activation contributes to SOX2 stabilization, Western blot analysis was first carried out to demonstrate that higher REST and SOX2 expression in ONS76 and DAOY cells is associated with increased p-AKT levels compared to the low-REST/low-SOX2 UW228 and UW426 cells (Fig. 3D). Next, we showed that *REST* knockdown in ONS76 and DAOY cells and its overexpression UW228 and UW426 promoted a reduction and increase in p-AKT levels (Figs. 3E and 3F). Then, the sensitivity of MB cell lines to the ATP-competitive inhibitor of AKT- Capivasertib was investigated. As seen in Fig. 3G, MTT assays showed that DAOY and ONS76 cells with high endogenous p-AKT were more sensitive to Capivasertib treatment (IC50 = 0.4 μM and 1.2μM, respectively) compared to UW228 and UW426 (IC50 = 12.3 μM and >20 μM, respectively) (Fig. 3G). Similar results were obtained with two other AKT inhibitors-Ipatasertib and Vevorisertib (Fig. S2)

To demonstrate that SOX2 stabilization in REST-driven SHH-MBs requires AKT activity, DAOY and ONS76 cells were treated with Capivasertib and the allosteric inhibitor MK2206, followed by Western blot analysis. Capivasertib treatment increased p-AKT levels as expected, but inhibited AKT kinase activity as evidenced by reduced phosphorylation of a known AKT target - GSK3β (Fig. 3H). MK2206 treatment also led to AKT inactivation, as suggested by the reduction in phosphorylation of both AKT and GSK3β (Fig. 3H). Importantly, AKT inactivation by both inhibitors caused a marked reduction in SOX2 levels in ONS76 and DAOY cells (Fig. 3H-I). Further, co-immunoprecipitation assays using anti-UBR5 antibody showed increased interaction between UBR5 and SOX2 upon treatment with AKT inhibitors (Fig. 3I). Collectively, these findings suggest that REST elevation drives AKT-mediated stabilization of SOX2 in SHH-MB cells.

### REST elevation upregulates Midkine-dependent cell-cell communication

Cell-cell communication is frequently altered in cancers, allowing tumor cells to override normal regulatory signals and reprogram their microenvironment to support tumor growth, metastasis, and resistance to therapy [45]. REST is known to control cell-cell communication for normal neuronal homeostasis through secreted messengers such as growth factors (e.g., BDNF [46]), cytokines (e.g., IL-1β [47]), and neuropeptides/hormones (e.g., CART [48] and VGF [49]). Whether REST influences cell-cell communication in tumors is not known. To investigate this possibility, we performed CellChat analysis to first show an increase in the number and strength of intercellular interactions within the malignant cell subclusters in high-REST compared to low-REST samples (191 vs. 143, and 4.347 vs. 2.758, respectively) (Figs. 4A and 4B). These data indicate that REST enhances the complexity and robustness of paracrine and autocrine communication between tumor cells, prompting further investigation into the input and output signaling patterns between high- and low-REST subclusters.

**Figure 4:**
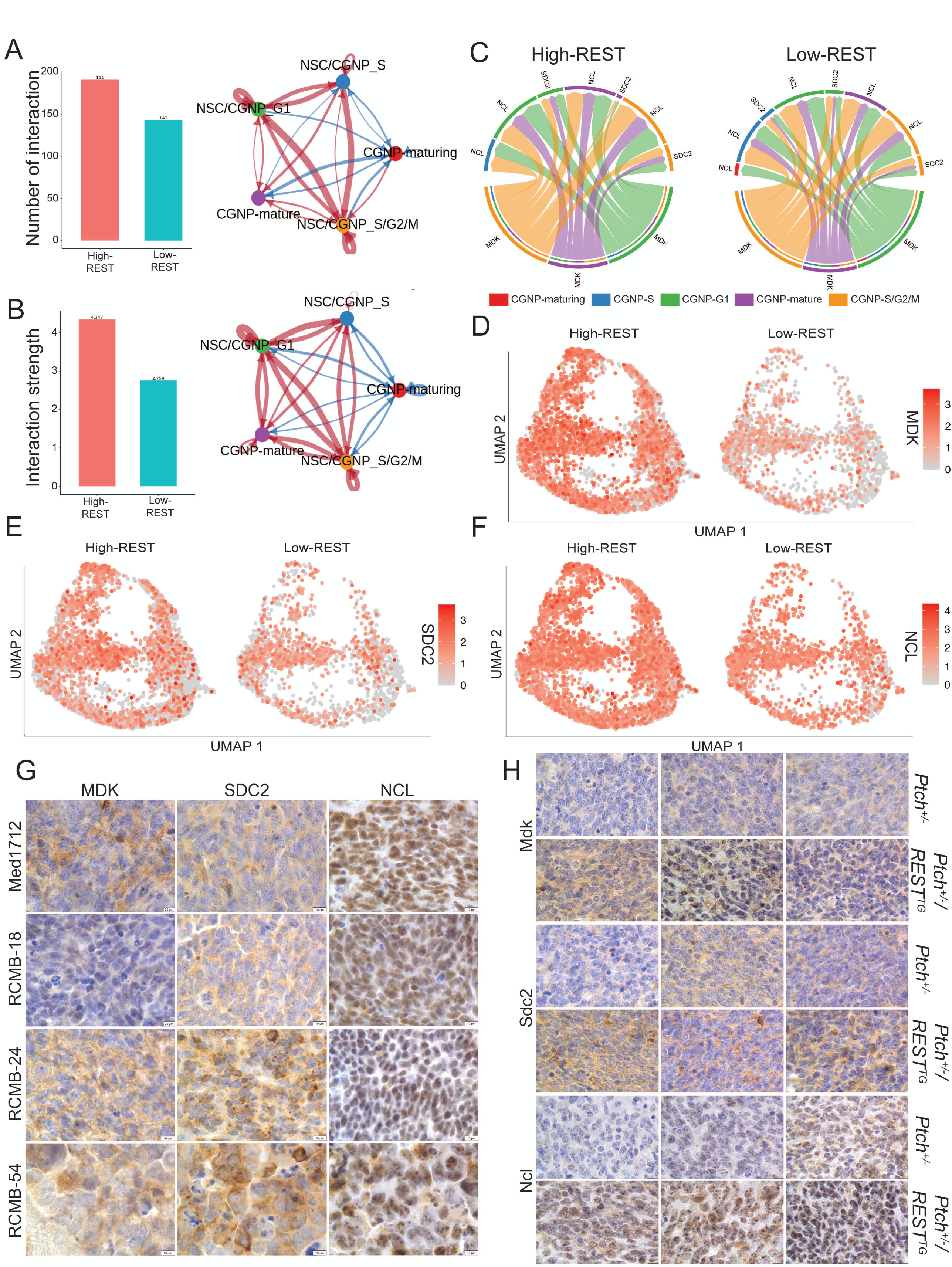
REST elevation is associated with increased MDK-mediated cell-cell communication in SHH-MBs. Bar graphs and circle plots depicting changes in the number **(A)** and strength **(B)** of inter- and intra-cluster cell-cell interactions in high-REST relative to low-REST CGNP clusters in SHH-MBs. **(C)** Circle plots show ligand-receptor interactions among CGNP subclusters in high- and low-REST SHH-MB samples. Colors denote CGNP cell states, and arcs show the strength and direction of MDK-dependent signaling interactions. **(D-F)** UMAP plot showing the expression of *MDK*, *SDC2* and *NCL* in the CGNP clusters from 1F. IHC to show MDK, SDC2, and NCL levels in tumor sections (**G**) from mice with SHH-MB PDOX and (**H**) *Ptch^+/−^* and *Ptch^+/−^/REST^TG^* animals (n = 3) (1000X magnification).

Input-output signaling analysis between subclusters revealed multiple ligand-receptor interactions that were altered in high-REST compared to low-REST subclusters. Some of these included Midkine (MDK)-Syndecan (SDC2) / MDK-Nucleolin (NCL), Neural Cell Adhesion Molecule 1 (NCAM1)-NCAM1 / NCAM1-fibroblast growth factor receptor 1 (FGFR1), Cadherin 2 (CDH2)-CDH2 and Cell Adhesion Molecule 1 (CADM1)-CADM1 (Fig. S3A-E). We focused on MDK-dependent signaling as a candidate for further studies in REST-driven SHH-MBs because of studies in lung adenocarcinoma, breast cancer, colorectal cancer and esophageal squamous cell carcinoma implicating this interaction in tumor progression [50–54]. Spatial and single-cell transcriptomic analyses of MBs have also shown elevated levels of MDK-dependent signaling interaction in high-risk tumors [28, 29]. However, mechanisms underlying increased MDK-dependent signaling are not known. As an initial step towards determining whether REST regulates ligand-receptor interactions in SHH-MBs, we showed increased MDK-SDC2 communication in high-REST tumors, with enhanced signaling among CGNP-G1, CGNP-S/G2/M, and CGNP-Mature clusters (Figs. S3A-C and 4C). MDK-NCL communication was similarly elevated in high-REST samples, with increased interactions among these same CGNP subclusters, including enhanced signaling of CGNP-G1, CGNP-S/G2/M with CGNP-G1 population (Figs. S3A–C and 4C). However, CGNP-S and CGNP-Maturing subclusters did not exhibit MDK-ligand associated outgoing interactions (Figs S2D-E and 4C). Chord plots also determined strong outgoing and incoming MDK interactions in CGNP-G1, CGNP-S/G2M, and CGNP-Mature populations in high-REST SHH-MB subclusters, in contrast to weaker and less frequent communications in the low-REST samples (Fig. 4C). Consistent with the CellChat analysis, UMAP expression plots identified markedly increased expression of MDK and its receptors, NCL and SDC2, in high-REST subclusters compared to low-REST subclusters (Fig. 4D-F). IHC staining of REST-expressing SHH-MB PDOX tumors also showed strong MDK staining within the extracellular space, while SDC2 localized predominantly to the cellular periphery in two samples (RCMB-24 and RCMB-54). The NCL showed a strong staining in all the four samples (Fig. 4G). Tumor-bearing brain sections from *Ptch^+/−^/REST^TG^* and Ptch^+/−^ mice confirmed a Rest-dependent elevation of Mdk and Sdc2. Interestingly, Ncl localization in tumors from Ptch^+/−^ mice was exclusively nucleolar, whereas cytoplasmic and membrane staining of Ncl was observed in *Ptch^+/−^/REST^TG^* tumor sections (Fig. 4H).

### The REST-SOX2 axis controls MDK and SDC2 expression

To understand if REST controls *MDK*, *SDC2* and *NCL* gene expression, SHH-MB patient transcriptomic datasets were analyzed to reveal a significant positive correlation between *REST* and *NCL* (r=0.2098; p=0.0016) (Figs. 5A), but not with *SDC2* or *MDK* (Fig. S4A). However, *SOX2* showed a strong positive correlation with both *MDK* (r=0.3886; p<0.0001) and *SDC2* (r=0.2694; p<0.0001) (Fig. 5B), but not with *NCL* (Fig. S4B). While NCL protein was expressed in DAOY, ONS76, UW228 and UW426 cells, SDC2 and MDK expression were higher in DAOY and ONS76 cells, which also have higher levels of REST (Fig. 5C). *REST* knockdown in ONS76 and DAOY cells caused a reduction in SDC2 and MDK levels, and conversely *REST* overexpression promoted an increase in the levels of these proteins in UW228 and UW426 cells (Figs. 5D and E). NCL levels were unaffected by changes in REST levels (Figs. 5D and E). The positive correlation of SOX2 with MDK and SDC2 (Figs. 5A-B) led us to investigate the requirement for SOX2 in REST-dependent changes in MDK and SDC2 levels. *SOX2* knockdown resulted in a decreased level of MDK, SDC2, and p-AKT in ONS76 and DAOY cells (Fig. 5F). Since *SOX2* knockdown promoted a significant decrease in *SDC2* and *MDK* mRNA expression and SOX2 showed significant binding to the promoter of *MDK* and *SDC2* in DAOY and ONS76 cells and intronic regions of *MDK* (ONS76), it indicates REST-dependent changes in MDK and SDC2 involve SOX2-mediated transcriptional regulation (Figs. 5G-J). Finally, treatment of DAOY and ONS76 cells with Capivasertib or MK2206 reduced MDK and SDC2 levels (Fig. 5K). Collectively, these findings support a model in which REST controls SOX2 stability through AKT-mediated interference of SOX2 degradation and SOX2 directly activates *MDK* and *SDC2* transcription in SHH-MB cells (Fig. 6). MDK-SDC2 signaling is known to activate PI3K-AKT pathway [55] indicating a feed forward loop that links REST-AKT-SOX2-MDK/SDC2 (Fig. 6).

**Figure 5:**
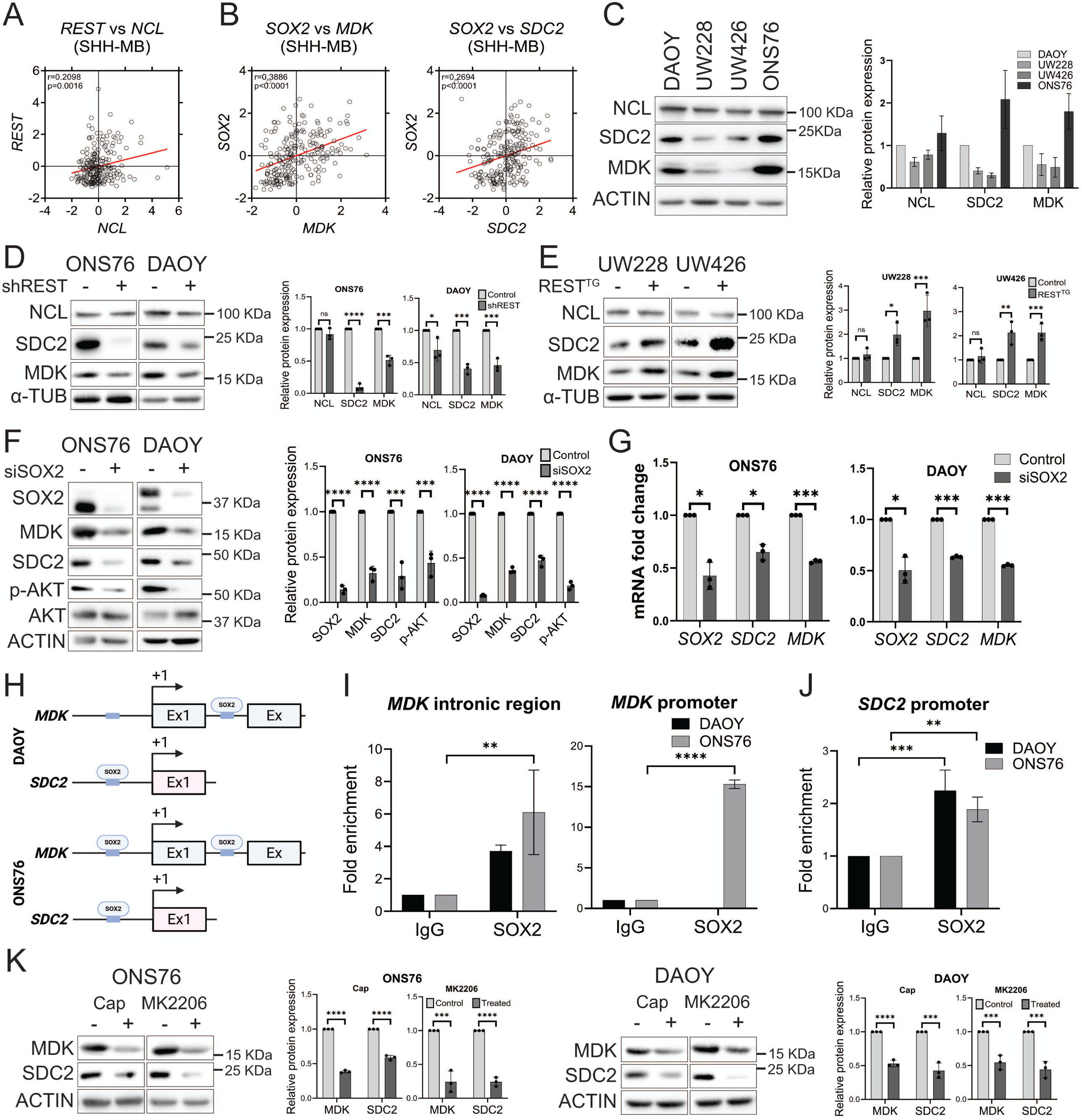
SOX2 controls *MDK* and *SDC2* expression in SHH-MB. Scatter plot to show correlation between **(A)** *REST* and *NCL* and **(B)** *SOX2* with *MDK* and *SDC2* genes in GSE85217 microarray dataset (n = 223). **(C)** Western blot to show basal levels of NCL, SDC2, and MDK in DAOY, ONS76, UW228 and UW426 cells and changes in their level following **(D)** REST knockdown and **(E)** its overexpression in ONS76/DAOY and UW228/UW426 cells, respectively. (**F**) Western blot to show changes in protein levels of SOX2, MDK, SDC2, pAKT (Ser473) and total AKT in ONS76 and DAOY cells following *SOX2* knockdown using specific esiRNA. **(G)** qPCR analyses to show reduction in *SOX2*, *MDK* and *SDC2* transcript following SOX2 knockdown using specific esiRNA in ONS76 and DAOY cells. **(H)** Location of SOX2 binding sites in *MDK* and *SDC2* genes. ChIP-qPCR analysis demonstrates SOX2 occupancy at **(I)** *MDK* intron and promoter-binding sites and **(J)** *SDC2* promoter in DAOY and ONS76 cells. For qPCR and ChIP-qPCR, statistics are presented for three independent experimental replicates as the means ± SDs. *p < 0.05, **p < 0.01, ***p < 0.001 and ****p < 0.0001 by Student’s t test. **(K)** Western blot analysis shows changes in MDK and SDC2 protein levels following treatment with Capivasertib and MK2206 in ONS76 and DAOY cells. Actin was used as a loading control. For all Western blot data, statistics are presented for three independent biological replicates as the means ± SDs. ns = non-significant, *p < 0.05, **p < 0.01, ***p < 0.001 and ****p < 0.0001 by Student’s t test.

**Figure 6.**
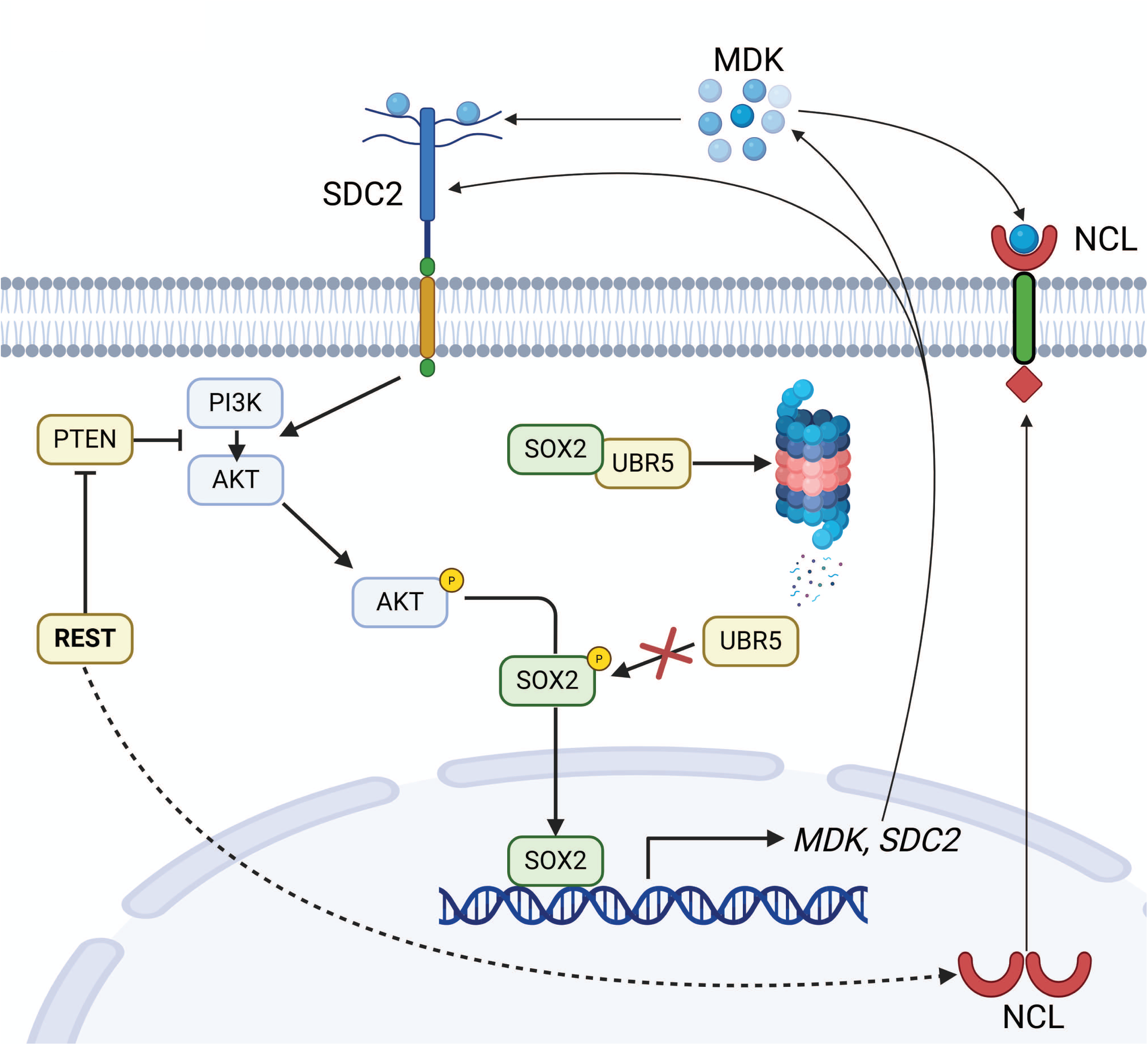
REST-driven AKT signaling stabilizes SOX2 and activates an MDK-SDC2/NCL feed-forward loop. REST negatively regulates PTEN expression, relieving inhibition of the PI3K-AKT pathway and leading to AKT activation. Activated AKT stabilizes SOX2 by preventing UBR5-mediated ubiquitination and proteasomal degradation. Stabilized SOX2 transcriptionally upregulates the expression of *MDK* and *SDC2*. In parallel, REST promotes the translocation of NCL from the nucleus to the cell membrane through an as-yet-undefined mechanism. Secreted MDK subsequently engages SDC2 and NCL at the cell surface, further activating PI3K-AKT signaling and reinforcing a feed-forward signaling loop.

## Discussion

Nearly 30% of MB patients experience relapses and insensitivity to standard of care [14, 56]. A growing body of evidence suggests this treatment insensitivity arises from both the acquisition of new genetic alterations following therapy and the persistence of a quiescent cancer stem-like compartment that survives initial treatment [13, 57]. In SHH-MB, this subpopulation comprises of neural precursor-like tumor stem cells marked by high expression of SOX2 and OLIG2, which originate from the CGNP lineage - cells of origin of SHH-MBs [58, 59]. Unlike the rapidly proliferating bulk tumor with hyperactive SHH signaling, SOX2⁺/OLIG2⁺ precursor cells maintain a more quiescent state, display enhanced DNA repair capacity, and can evade cytotoxic chemotherapy and radiation [13, 57, 60]. Interestingly, SOX2^+^/REST^+^ precursor cells exhibit a higher proliferative capacity. This raises important questions about the cellular niche in the SOX2⁺/OLIG2⁺ and SOX2^+^/REST^+^ contexts. Nevertheless, survival of both these cell types has the potential to drive tumor regrowth, recurrence and poor outcomes in patients with SHH-MBs [60].

Regulation of stem cell identity is a complex process and involves integrated input from transcription factors, chromatin remodeling and accessibility, metabolic and microenvironmental cues [61–63]. In SHH-MBs, signaling is primarily driven by the transcription factors GLI1 and GLI2, where GLI2 acts as the upstream pioneer activator and GLI1 serves as an amplifier of pathway output [64, 65]. Notably, GLI2 and SOX2 form a positive regulatory feedback loop, where GLI2 binds the *SOX2* promoter-proximal region to enhance transcription [42, 66].

Here, we show that elevated REST expression in SHH-MBs is associated with enrichment of stem cell-related pathways, consistent with literature demonstrating REST’s central role in maintaining an immature cell state through repression of neuronal differentiation programs [22, 23]. REST is known to restrain cell-cycle exit in CGNPs, promoting a proliferative progenitor state [41]. The preferential enrichment of SOX2⁺ cells among cycling CGNPs with elevated REST expression suggests that REST not only sustains proliferation but may also expand or preserve a more primitive, neural stem-like compartment within SHH-MBs.

In the current study, we identified a shift toward higher *GLI2* expression in high-REST SHH-MBs, providing a potential mechanistic basis for increased *SOX2* expression. Thus, REST likely promotes neural stem-like states while simultaneously biasing SHH pathway activity toward GLI2 dependence. Although a direct role for REST in regulating *GLI2* transcription cannot be excluded, a more plausible indirect mechanism may involve MYC, which has been shown to cooperate preferentially with GLI2 to sustain SHH transcriptional output in stem-like contexts [67]. Through transcriptional amplification, modulation of chromatin accessibility at GLI2 target loci, and repression of differentiation-associated negative regulators of SHH signaling, MYC could reinforce GLI2-driven programs in high-REST tumors. As stated above, other groups have shown a role for OLIG2 in regulating stem cell fate transitions and promoting neoplastic progenitor generation in SHH-MBs [57, 60]. However, our analyses did not identify OLIG2 as a key driver of SOX2 positivity in REST-driven SHH-MBs. A potential role for REST in remodeling the SOX2 regulome in this setting remains an important area for future investigation.

These transcriptional and signaling biases prompted us to investigate whether REST also regulates SOX2 abundance through post-translational mechanisms. We identify a post-transcriptional mechanism in human SHH-MB whereby AKT blocks UBR5-mediated proteasomal degradation of SOX2, leading to its stabilization [44, 68, 69]. REST elevation is associated with downregulation of *Pten* expression and AKT signaling hyperactivation [20], and is consistent with our finding that REST elevation interferes with UBR5/SOX2 interaction and inhibition of AKT activity facilitate SOX2 destabilization. These data suggest that AKT inhibition can be used to sensitize REST-driven MBs to standard of care or tested in combination with inhibitors of SHH signaling.

Cell-cell communication within tumors, particularly among cancer stem cells (CSCs) and their supporting niche, is a critical determinant of therapeutic resistance and relapse [70, 71]. Such interactions can reinforce CSC survival through paracrine and autocrine signaling pathways, direct cell-cell contacts, and the exchange of regulatory factors, to collectively promoting quiescence and resistance to cytotoxic therapies [72, 73]. Here, we identify a previously unrecognized role for malignant cell-intrinsic communication via the MDK-SDC2/NCL axis in amplifying AKT signaling and stabilizing SOX2 in REST-driven SHH-MBs. MDK signaling activates PI3K-AKT-mTOR and MAPK-ERK pathways, which support CSC maintenance, survival, and EMT-like programs [74, 75]. In addition, MDK is implicated in protecting tumor cells from apoptosis and promoting the expression of drug-efflux transporters, including ABC family members and P-glycoprotein, suggesting a broader role in mediating therapy resistance that warrants further investigation in REST-high SHH-MBs [76, 77].

While elevated MDK signaling is reported in human MBs [28, 29], the regulatory mechanisms driving its aberrant expression and activation have remained undefined. To our knowledge, this study is the first to identify REST and SOX2 as upstream regulators of MDK and its receptors SDC2 and NCL in SHH-MBs. Our data suggests that *MDK* and *SDC2* are transcriptional targets of SOX2, whereas subcellular localization of NCL rather than its expression is regulated by REST. The pronounced REST-dependent redistribution of NCL from the nucleolus to the cytoplasmic/pericellular compartment raises the intriguing possibility that REST functionally reprograms NCL to serve as a signaling receptor for MDK to reinforce AKT activation. Although the precise molecular mechanisms underlying this re-localization remain to be defined, these findings position the MDK-SDC2/NCL axis as a critical signaling node downstream of REST and SOX2 in SHH-MB stem-like cells.

Thus, our work is among the first to identify a role for REST in the post-translational regulation of SOX2 in SHH-MBs and in the control of MDK-SDC2/NCL signaling upstream of AKT activation. Our findings support the existence of a positive feedback loop in which REST-driven AKT signaling stabilizes SOX2, while SOX2 in turn upregulates MDK and SDC2 to further potentiate AKT pathway activity (Fig. 6). This REST-AKT-SOX2-MDK/SDC2 circuitry defines a previously unrecognized signaling axis that sustains stem-like states in high-REST SHH-MBs and represents a potential therapeutic vulnerability. Disruption of this pathway, either through AKT inhibition or targeting MDK signaling with small-molecule inhibitors such as HBS-101 [78] or neutralizing antibodies [79], merits investigation in future pre-clinical studies.

## Acknowledgements

The work was supported by grants from Addi’s Faith Foundation to VG. We thank Dr. Rong-Hua Tao for helpful discussions.

## Author Contribution

AS, LX, and VG designed experiments and study. AS, LG, JY, DC, YY, and SM performed experiments. AS, JS, and VG analyzed the data. AS, LG, and VG wrote the manuscript. VG provided funding support.

---

**Figure S1:**
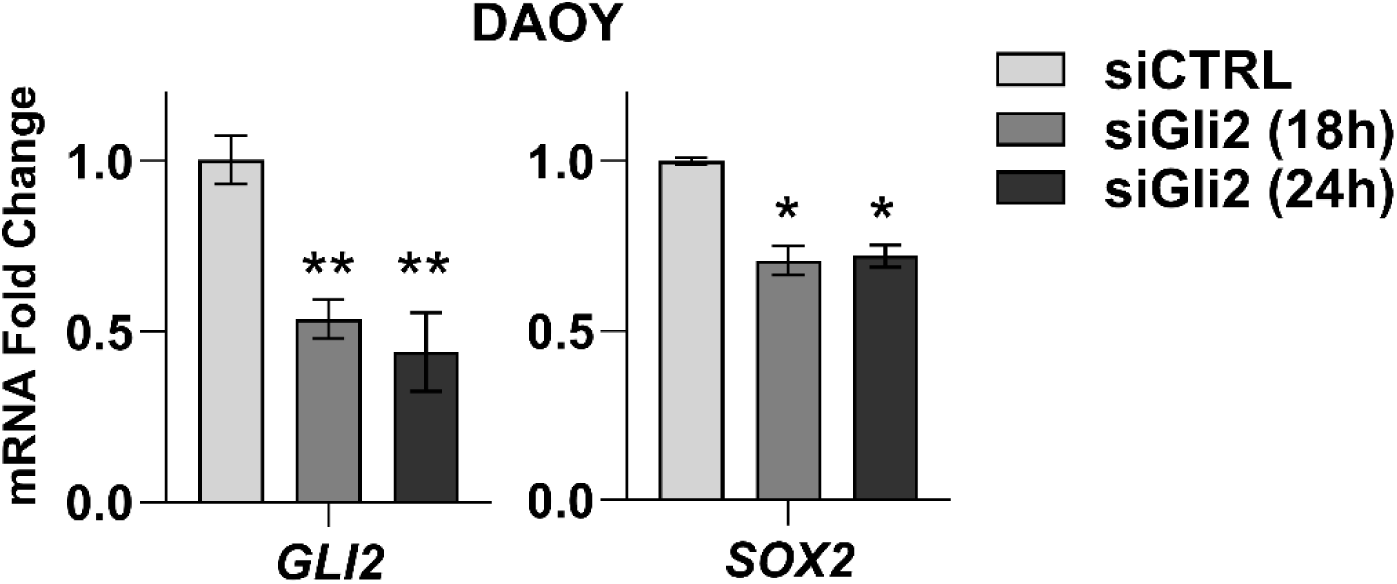
qPCR analysis showing reduced *GLI2* and *SOX2* transcript levels following siRNA-mediated GLI2 knockdown for 18 and 24 h in DAOY cells.

**Figure S2:**
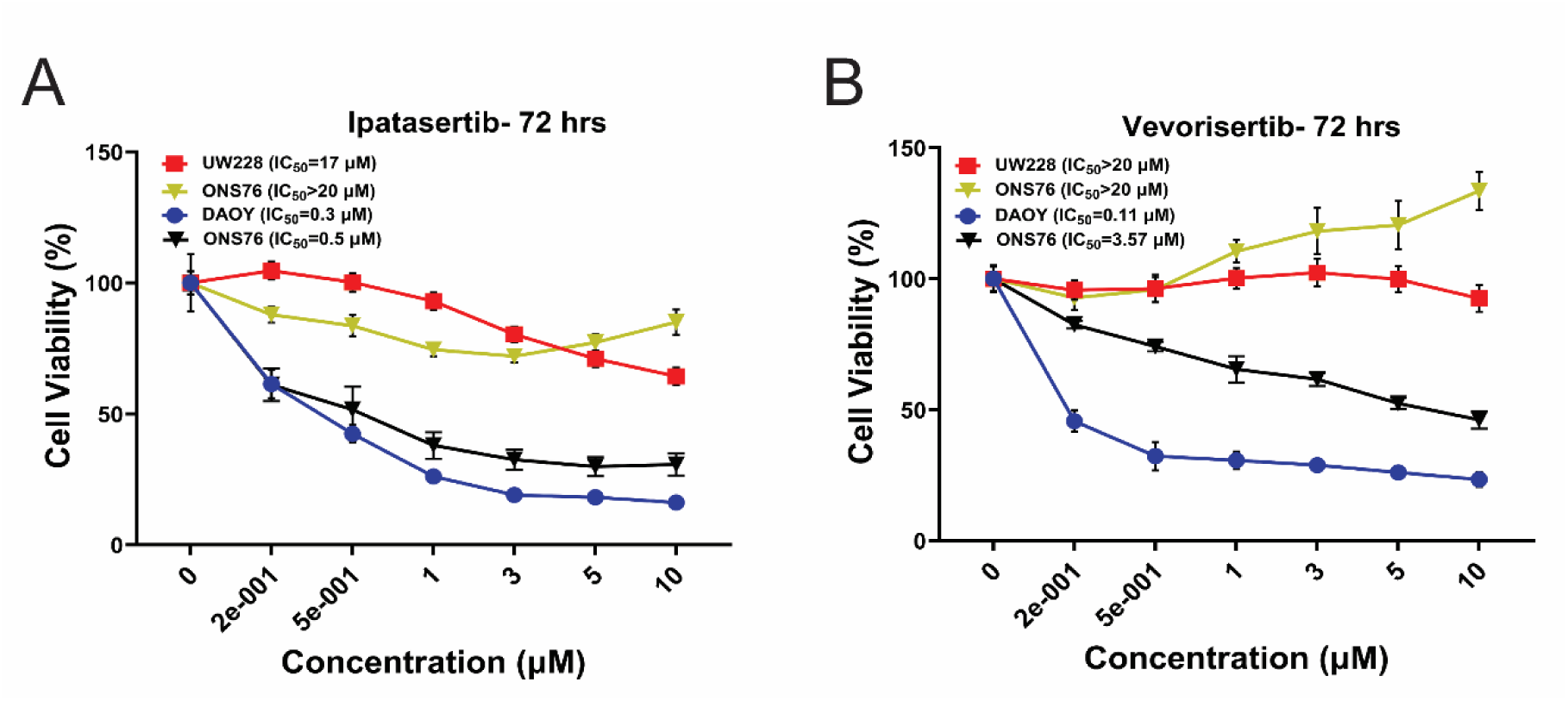
Dose–response curves for SHH-MB cell lines (DAOY, UW228, UW426, and ONS76) following 72-hour treatment with increasing concentrations of Ipatasertib **(A)** and Vevorisertib **(B)**. IC50 values for each cell line are shown in the inset. All experiments were done in triplicate.

**Figure S3:**
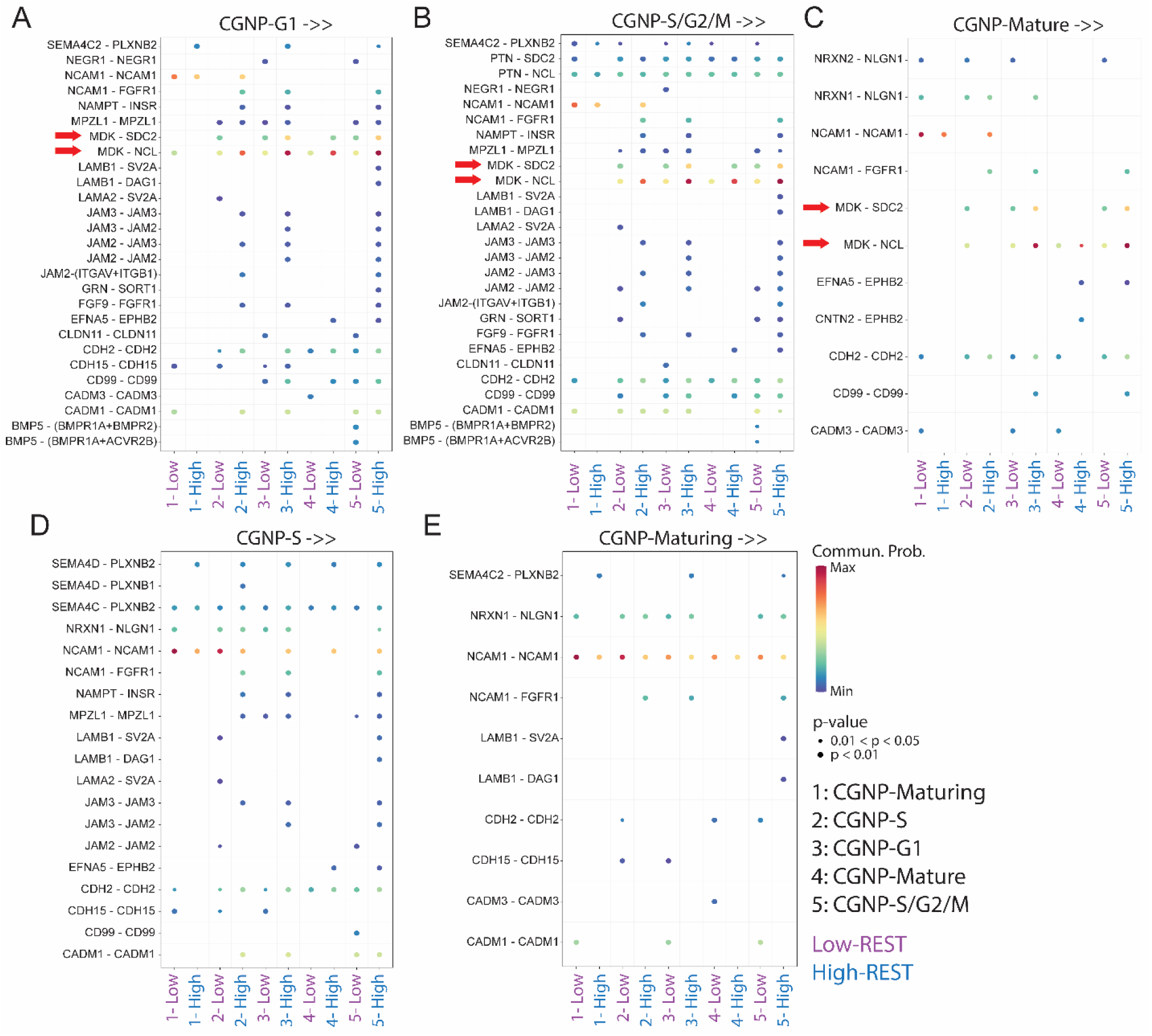
Bubble plot depicting ligand–receptor interactions between ligands expressed in CGNP-G1 **(A)**, CGNP-S/G2M **(B)**, CGNP-Mature **(C)**, CGNP-S **(D)**, and CGNP-Maturing **(E)** clusters and receptors expressed in other cell clusters (clusters 1–5) in high- and low-REST populations. Dot size represents the fraction of cells expressing the indicated gene, and dot color denotes the average gene expression level within each lineage. MDK-mediated signaling interactions are highlighted by a red arrow.

**Figure S4:**
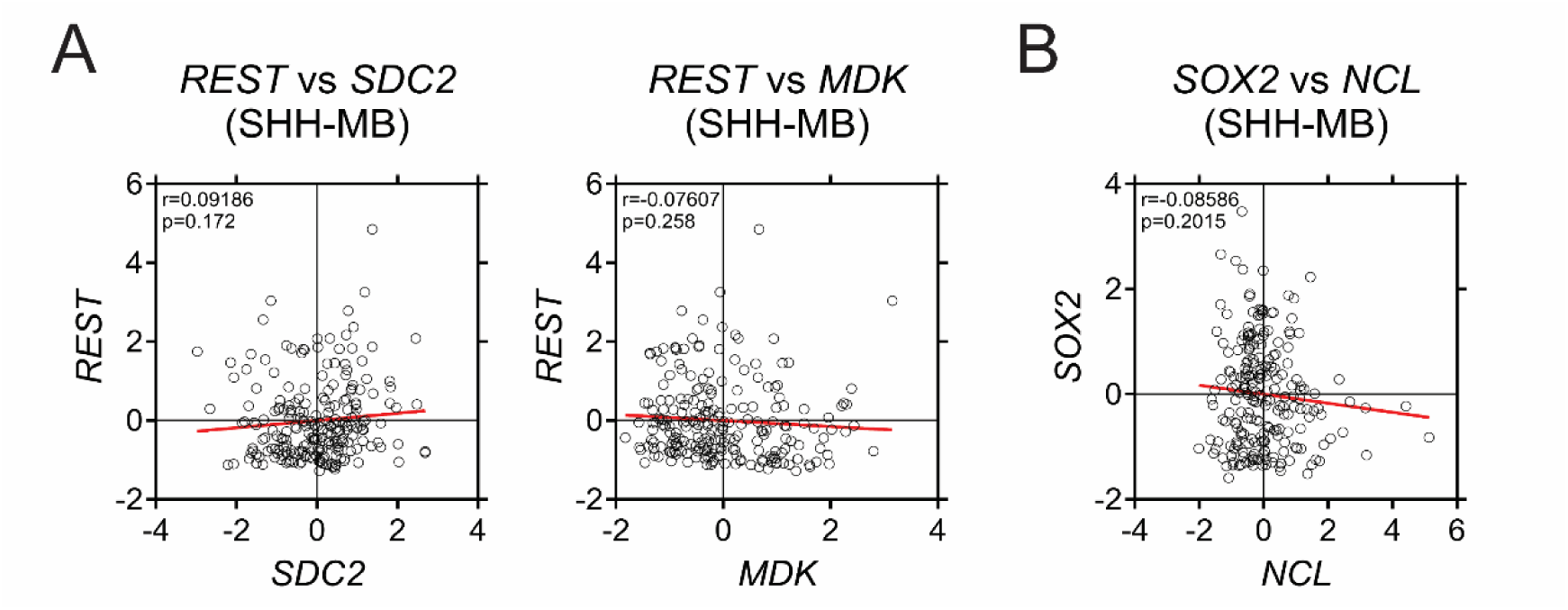
Scatter plot showing the correlation of mRNA expression of *REST* with *SDC2* and *MDK* (A) and *SOX2* with *NCL* (B) in GSE85217 dataset (n = 223).

**Table S1:** List of antibodies.

| **Antibody** | **Company** | **Catalogue number** |
| --- | --- | --- |
| **Western blot** |  |  |
| REST | Sigma-Aldrich | 07-579 |
| SOX2 | Abcam | ab97959 |
| Akt | Cell Signaling Technology | 9272 |
| Phospho-Akt (Ser473) | Cell Signaling Technology | 3787 |
| GSK-3β | Cell Signaling Technology | 9315 |
| Phospho-GSK-3β (Ser9) | Cell Signaling Technology | 9323 |
| UBR5 | Cell Signaling Technology | 65344 |
| Midkine | Thermo Fisher Scientific | 11009-1-AP |
| Syndecan2 | Thermo Fisher Scientific | 36-6200 |
| Nucleolin | Cell Signaling Technology | 14574 |
| α-Tubulin | Cell Signaling Technology | 9099 |
| Histone H3 | Thermo Fisher Scientific | PA5-16183 |
| Actin | Millipore Sigma | A1978 |
| **Immunohistochemistry** |  |  |
| REST | Abcam | ab202962 |
| GLI2 | Thermo Fisher Scientific | 18989-1-AP |
| Ki67 | Abcam | ab15580 |
| TUBB3 | Abcam | ab18207 |
| SOX2 | Abcam | ab97959 |
| PTEN | Cell Signaling Technology | 9188 |
| Phospho-Akt (Ser473) | Cell Signaling Technology | 3787 |
| Midkine | Thermo Fisher Scientific | 11009-1-AP |
| Syndecan2 | Thermo Fisher Scientific | 36-6200 |
| Nucleolin | Cell Signaling Technology | 14574 |
| **Co-immunoprecipitation** |  |  |
| UBR5 | Cell Signaling Technology | 65344 |
| **Chromatin-immunoprecipitation** |  |  |
| SOX2 | Thermo Fisher Scientific | MA1-014 |

**Table S2:**
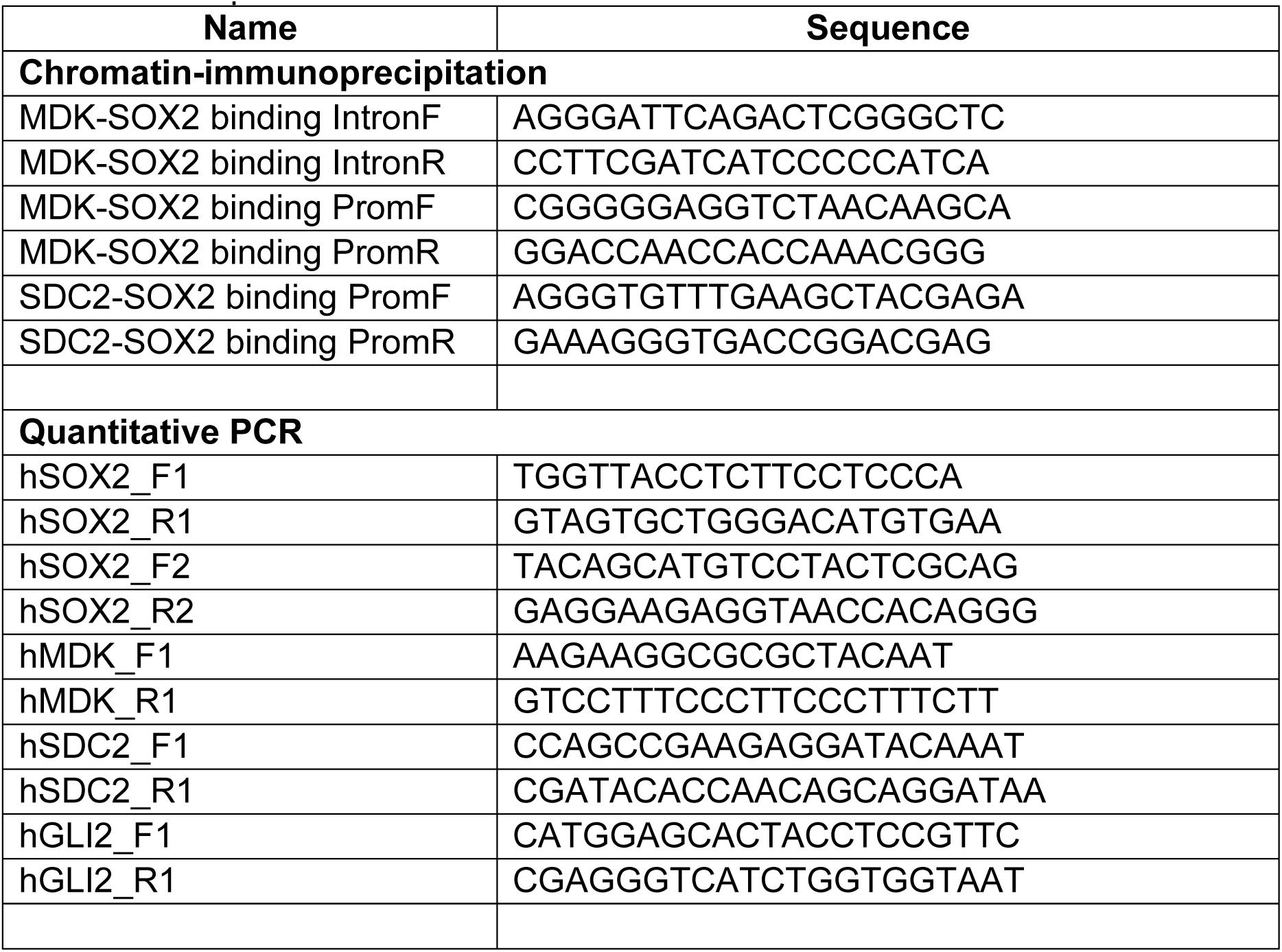
List of primers.

